# Computational analysis of B cell receptor repertoires in COVID-19 patients using deep embedded representations of protein sequences

**DOI:** 10.1101/2021.08.02.454701

**Authors:** Inyoung Kim, Sang Yoon Byun, Sangyeup Kim, Sangyoon Choi, Jinsung Noh, Junho Chung, Byung Gee Kim

## Abstract

Analyzing B cell receptor (BCR) repertoires is immensely useful in evaluating one’s immunological status. Conventionally, repertoire analysis methods have focused on comprehensive assessments of clonal compositions, including V(D)J segment usage, nucleotide insertions/deletions, and amino acid distributions. Here, we introduce a novel computational approach that applies deep-learning-based protein embedding techniques to analyze BCR repertoires. By selecting the most frequently occurring BCR sequences in a given repertoire and computing the sum of the vector representations of these sequences, we represent an entire repertoire as a 100-dimensional vector and eventually as a single data point in vector space. We demonstrate that this new approach enables us to not only accurately cluster BCR repertoires of coronavirus disease 2019 (COVID-19) patients and healthy subjects but also efficiently track minute changes in immune status over time as patients undergo treatment. Furthermore, using the distributed representations, we successfully trained an XGBoost classification model that achieved a mean accuracy rate of over 87% given a repertoire of CDR3 sequences.

## Introduction

Analyzing and deciphering biological sequences plays a critical role in gaining a deeper understanding of biological systems. Recent major advances in artificial intelligence have sparked ample interest in adopting natural language processing (NLP) models to extract hidden insights from biological sequences [1]. By reinterpreting protein sequences as sentences and k-mers in these sequences as words, researchers have succeeded in establishing computational methods to represent the language of life. One of the most widely used techniques in the field is Words2Vec, an efficient method to learn and compute word embeddings, which are essentially vectorized representations of words [2].

ProtVec was the first bioinformatics application to use such word-vector model to embed biological sequences—more specifically, amino acid sequences [3]. The representation, which treats amino acid 3-mers as discrete words and employs the skip-gram model architecture, has been rigorously trained on over 546,790 unique protein sequences from the Swiss-Prot database. Ultimately, ProtVec is highly efficient in transforming an amino acid sequence into a 100-dimensional vector.

B-cell receptors (BCRs) are transmembrane proteins on the surfaces of B-cells that are crucial in the production of antibodies that recognize and neutralize a myriad of foreign objects like antigens [4]. To bind to a wide range of antigens, BCRs most often undergo genetic recombination and somatic hypermutation (SHM), ultimately generating a more diversified pool of paratopes, also known as antigen-binding sites. Each paratope comprises six complementarity-determining regions (CDRs), including three each from the antibody’s heavy and light chains. Moreover, CDRs can be categorized into three distinct sections (CDR1, CDR2, and CDR3), where CDR3 is commonly deemed the most variable, and hence the most critical region for recognizing antigens. Therefore, this study will primarily focus on heavy-chain CDR3s when discussing CDRs.

A BCR repertoire refers to a diverse population of BCRs in a given individual or entity. Analyzing BCR repertoires is highly useful in determining and evaluating the overall condition of one’s immune system [5]. For example, a healthy individual will generally have a diverse repertoire of unique sequences to recognize a variety of antigens. A virus-infected individual will similarly have a wide variety of sequences, but they will also most likely have aberrant increases in the numbers of certain BCR sequences due to the activation of specific B-cells that bind to invading antigens [6]. BCR repertoire analysis provides a holistic assessment of one’s adaptive immune system as well as immune diversity.

Previous studies on BCR repertoires have primarily focused on classifying and clustering BCRs based on sequence motifs in an attempt to identify neutralizing antibodies against specific viruses [7–10]. One of the most common and conventional methods for studying BCRs in a wet lab is flow cytometry, which aims to identify antigen-specific B-cells by labeling antigens with fluorescent t ags. Microscopy and immunohistochemical staining are other common techniques used to detect particular antigens.

Recently, several computational methods have also been established. Schulteis, for example, employed the GLIPH2 algorithm to cluster T-cell receptor (TCR) sequences and the OLGA algorithm to compute CDR3 sequence similarities from TCR/BCR repertoires of coronavirus disease 2019 (COVID-19) patients [11]. Bashford-Rogers developed a method that utilizes network analysis to compare a wide range of BCR repertoires [12]. Sidhom very recently proposed DeepTCR, a deep learning model that enables enhanced classification of antigen-specific TCRs from human and murine datasets [13].

In this study, we present a novel computational approach for representing BCR repertoires using a deep-learning based protein embedding method. By utilizing ProtVec to embed all BCR amino acid sequences and summing the vectorized values of the most frequently occurring CDR3 sequences in a given repertoire, we represent individual BCR repertoires as single 100-dimensional vectors. Our representation shows promising results for clustering, classifying, and even tracking BCR repertoires of COVID-19 patients and healthy individuals in vector space. Compared to conventional techniques, this computational method provides an entirely new approach for researchers to characterize and analyze BCR repertoires.

## Methods

### Statistical information

Twenty-five studies, including 3 COVID-19 studies, were downloaded and used for analysis. In total, this study analyzed 106 COVID-19 patient repertoires and 349 healthy subject repertoires (later reduced to 322 due to a lack of sufficient unique CDR3 sequences in 27 healthy subjects).

### Embedding amino acid sequences using ProtVec

ProtVec regards amino acid 3-mers as ‘biological words’ and effectively embeds each word (over 9,048 words in total) into a 100-dimensional vector. As shown in Fig. 1, an original sequence is first split into three separate lists of nonoverlapping 3-mers. The 3-mers in these lists are then converted into vectors based on pretrained data available from the Harvard Dataverse [3]. Finally, by summing all the vector representations of the nonoverlapping 3-mers, a single 100-dimensional vector that represents a single protein sequence is obtained. The pretrained data file available from the Harvard Dataverse is a comma-separated values (CSV) file that includes 100 float values assigned to each of the 9,048 3-mers. The model was trained on 546,790 sequences from the Swiss-Prot database using the Skip-gram Word2Vec architecture. The CSV file can be used as a look-up table to search for 100-dimensional vectors based on a given 3-mer.

**Fig. 1.**
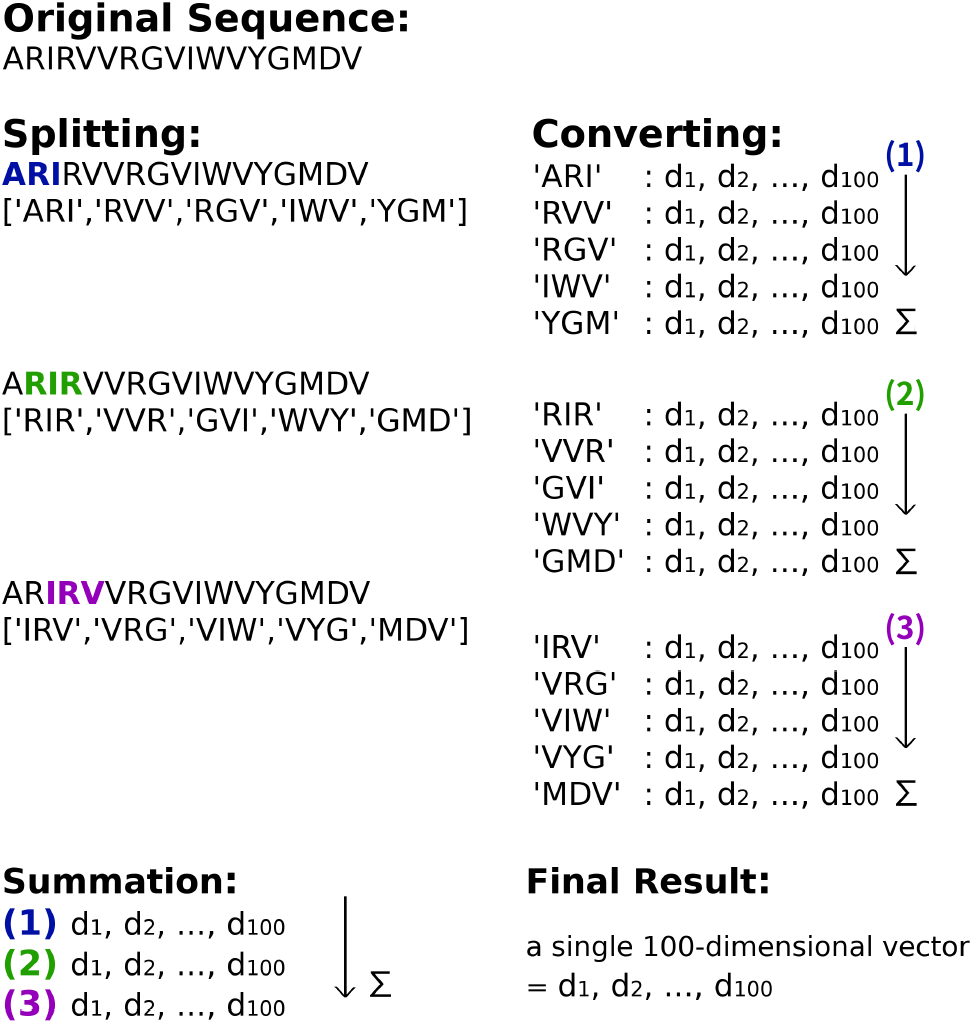
Embedding protein sequences using ProtVec. An illustration of embedding an amino acid sequence using ProtVec. This technique enables one to express a protein sequence as a single 100-dimensional vector while retaining its unique physical/chemical properties.

### Data acquisition and preprocessing

The raw data acquired from the Observed Antibody Space (OAS) BCR dataset included heavy and light chains as well as several different isotypes. We decided to use sequences from the immunoglobulin heavy G chain (IGHG) due to its strong association with severe acute respiratory syndrome coronavirus 2 (SARS-CoV-2) [7]. The raw data did not include read counts of unique CDR3 sequences, but did include read counts of unique BCR sequences. As a result, we added a preprocessing step that counted the number of unique CDR3 sequences in a given repertoire.

For the attributes used to search the OAS sequences, we set ‘Chain’ to ‘Heavy’, ‘Isotype’ to ‘IGHG’, ‘Disease’ to ‘SARS-COV-2’ or ‘None’, ‘BSource’ to ‘PBMC’, ‘Vaccine’ to ‘None’, and ‘Species’ to ‘human’. For the attributes used to search the OAS sequences, we set ‘Chain’ to ‘Heavy’, ‘Isotype’ to ‘IGHG’, ‘Disease’ to ‘SARS-COV-2’ or ‘None’, ‘BSource’ to ‘PBMC’, ‘Vaccine’ to ‘None’, and ‘Species’ to ‘Human’. Twenty-five studies, including 3 COVID-19 studies, were downloaded and used for analysis. In total, this study analyzed 106 COVID-19 patients and 349 healthy subjects (later reduced to 322 due to a lack of sufficient unique CDR3 sequences in 27 healthy subjects).

### Visualizing sequences and repertoires using t-SNE

Visualizing the embedded representations (sequences and/or repertoires) in a 2-dimensional vector space requires reducing the 100-dimensional ProtVec data to 2-dimensional x, y coordinates so that one can ultimately generate a scatter plot. We utilized the t-distributed stochastic neighbor embedding technique (t-SNE implementation from scikit learn) to reduce the dimensions. For all the t-SNE charts shown herein, we set the specific hyperparameters to ’perplexity=30.0’ and ’learning_rate=100’.

### Training binary classification models

For model training, we utilized the support vector machine (SVM) implementation from scikit learn library (version 0.22.2.post1) and XGB-Classifier from the XGBoost Python package (version 0.90). For the SVM model, which classified the two selected proteins families (Fig. 2), we used 1,528 embedded representations of amino acid sequences from each family (3,056 in total). For the XGBoost model, which classified COVID-19 patients and healthy subjects (Fig 4b), we used 106 embedded representations of BCR repertoires from each of the two classes (212 in total). We evaluated both models using 5-fold cross validation (StratifiedKFold implementation from scikit-learn) and receiver operating characteristic/area under the curve (ROC/AUC) computation (scikit-learn). All sequences and/or repertoire data that were involved in training, testing, and validation of the models were embedded using ProtVec.

**Fig. 2.**
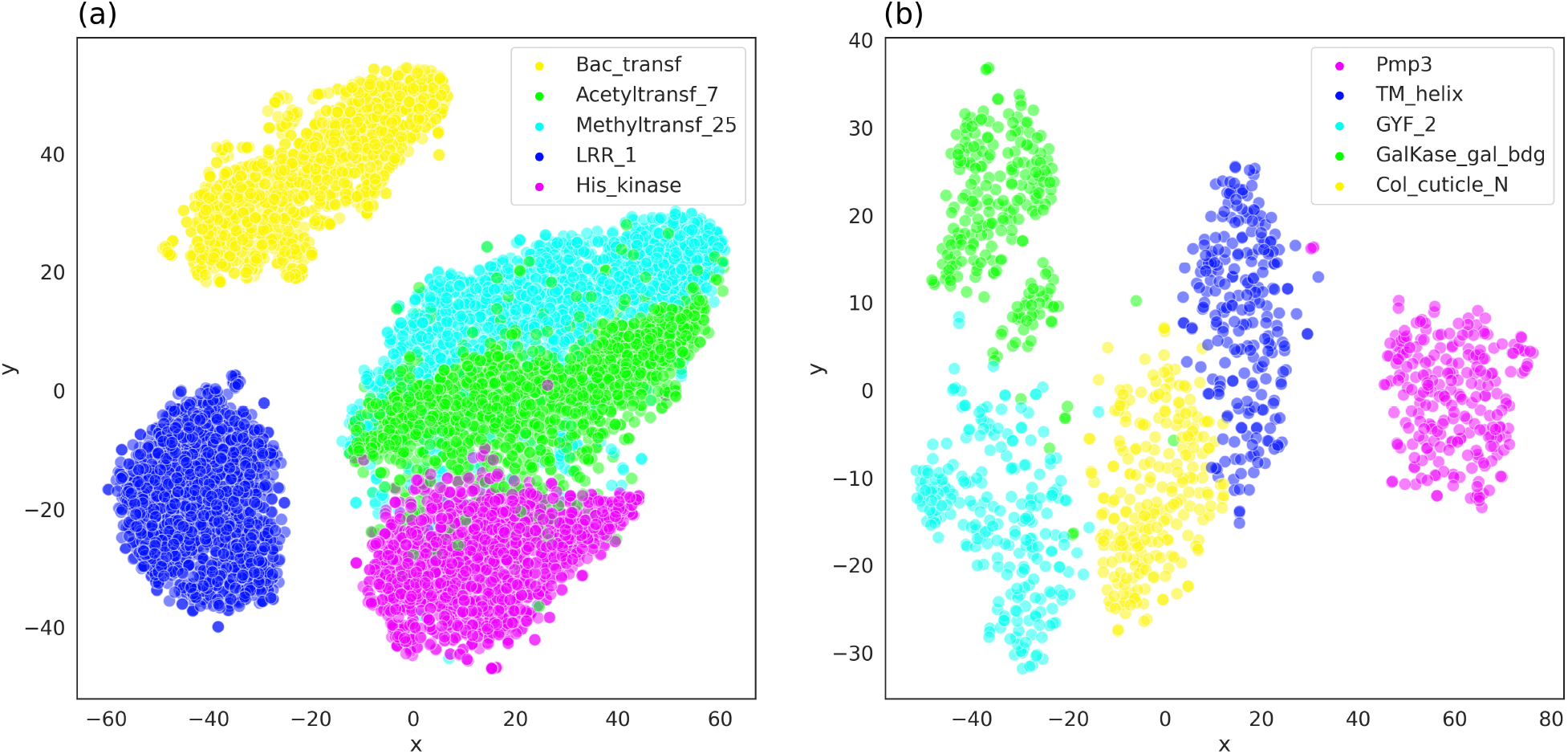
Visualization of embedded protein sequences. Result of embedding the acquired protein sequences and visualizing them on a 2-dimensional vector space using t-SNE. **(a).** Visualization of amino acid sequences from five protein families that had the largest number of sequences in the dataset. A total of 7,640 sequences (1,528 sequences from 5 families) were used. Detailed list of selected protein families, including their corresponding family ID, the number of sequences, and family accession numbers, is summarized in Table 1a. **(b).** Visualization of sequences from five similar (in terms of mean sequence length and standard deviation) protein families. A total of 1,210 sequences (242 sequences for 5 families) were used. Detailed information about the protein families is summarized in Table 1b.

## Results

### Assessment of the efficacy of ProtVec representation

To ensure that ProtVec can indeed accurately and precisely represent protein sequences, we initially assessed the efficacy of the embedding technique in a public dataset provided by Google AI [14]. Details on the specific protein families and the number of sequences used in the assessment are shown in Table 1. After embedding each of the protein sequences as 100-dimensional vectors, we used t-SNE to reduce their dimensions. We observed that the protein families of interest were successfully clustered into five distinct classes when plotted in 2-dimensional vector spaces (Fig. 2a and 2b). As shown in Fig. 2a, we found that two classes, Methyltransf_25 and Acetyletransf_7, overlapped with each other in the t-SNE chart, possibly hinting that t-SNE was unable to distinguish the two families.

**Table 1.** Protein families. Details regarding family ID, the number of CDR3 sequences, family accession numbers, and mean sequence lengths of the selected sequences from the Google AI public protein dataset.

| Family ID | No. of CDR3s | Family Accession |
| --- | --- | --- |
| Methyltransf_25 | 3,637 | PF13649.6 |
| LRR_1 | 1,927 | PF00560.33 |
| Acetyltransf_7 | 1,927 | PF13508.7 |
| His_kinase | 1,537 | PF06580.13 |
| Bac_transf | 1,528 | PF02397.16 |

| Family ID | No. of CDR3s | Accession | Mean Length |
| --- | --- | --- | --- |
| GalKase_gal_bdg | 448 | PF10509 | 49.82 |
| Col_cuticle_N | 242 | PF01484 | 49.41 |
| TM_helix | 247 | PF05552 | 49.38 |
| GYF_2 | 307 | PF14237 | 49.16 |
| Pmp3 | 286 | P87284 | 49.14 |

When we used SVM models to further analyze the two families, we obtained a classification model that successfully classified the two families with 97% mean accuracy and an AUC value of 0.99. Thus, we eventually came to the conclusion that the overlap was a result of some negligible limitation that may have occurred during the process of dimensionality reduction. Specifically, for the five protein families with similar lengths, we first examined the mean lengths of the protein sequences from each of the protein families and ultimately selected five families that had similar mean lengths (49 amino acids). We also searched for families that had low standard deviations to ensure that the sequences from the selected families did not vary widely in length. The plotted sequences from five protein families, which are presented in Fig. 2b, were similar in mean sequence length, and we observed five distinct clusters with minimal overlap.

### The distribution of frequencies of CDR3 read counts

Before closely analyzing the repertoires, we briefly explored the arrangements of few different repertoires to gain a better understanding of the distribution of CDR3 sequences. Fig. 3a shows the histogram of CDR3 amino acid sequences from a BCR repertoire of a COVID-19 patient (Subject-A_TP2; Kim et al.) and that of a healthy subject (700010756; Roskin et al.). In comparing the two histograms, we observed that the patient’s repertoire (purple) possessed more unique CDR3 sequences in general and many more CDR3 sequences with higher read counts than those from the healthy individual’s repertoire (green). While the healthy repertoire contained a sequence with a read count of 9,901, which was abnormally high, all other sequences in the repertoire had significantly lower read counts than the read counts in the patient repertoire (Table 2). Conversely, the highest read count in the patient’s repertoire was 2,631, and the succeeding four sequences also had read counts higher than 1,000, suggesting that sequences in the patient’s repertoire generally has much higher read counts.

**Fig. 3.**
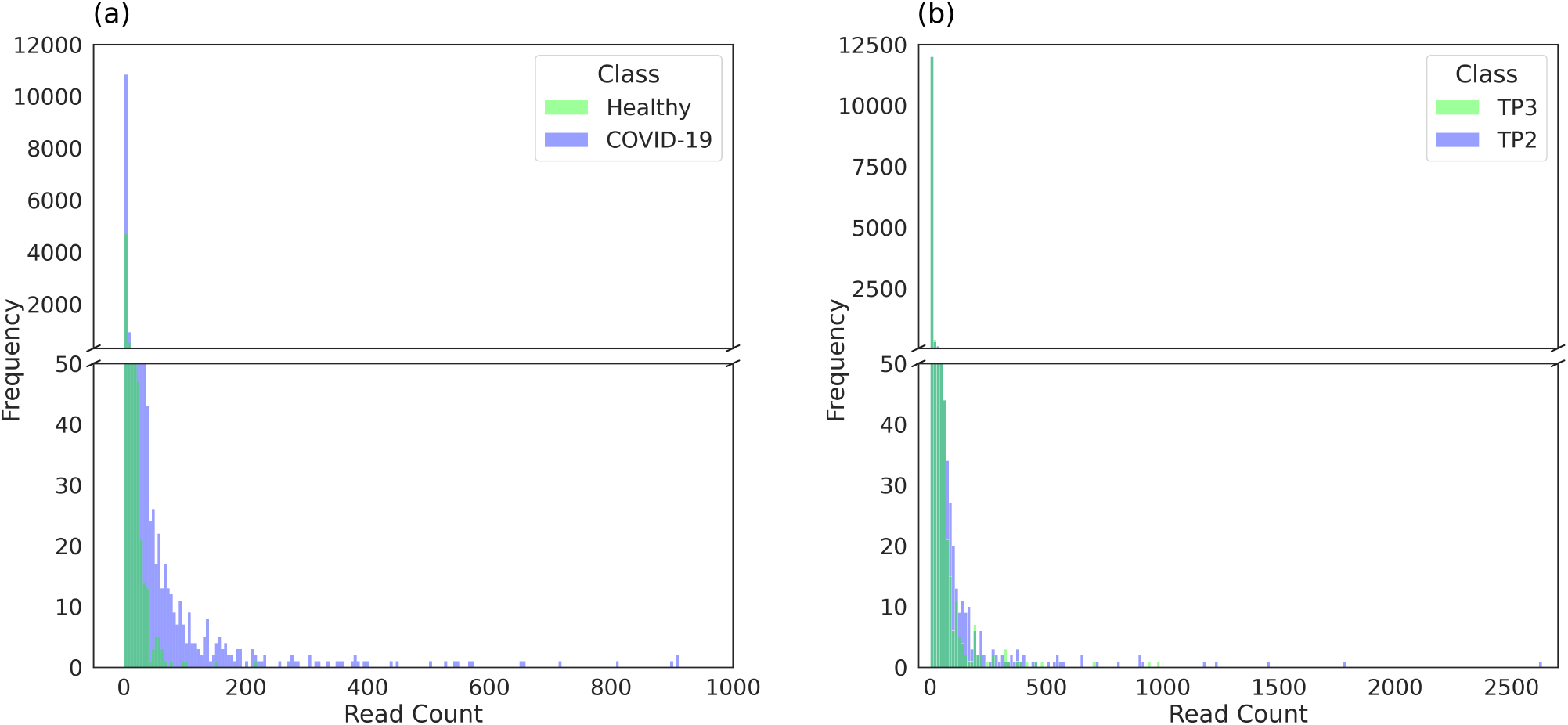
The distribution of CDR3 read counts. The distributions of frequencies of CDR3 sequence read counts between a patient BCR repertoire and a healthy repertoire. Read count (x-axis) represents the number of times a specific CDR3 sequence appears within a repertoire, and frequency (y-axis) represents the number of times the corresponding read count occurred. **(a)**. A histogram (bins = 2,000) of repertoires from a COVID-19 patient and a healthy subject. For the sake of clear visualization, we display only sequences with read counts up to 1,000. More details can be found in Table 2. **(b)**. A histogram (bins = 200) of repertoires from Subject A’s two time points (from the study by Kim et al.). TP2 is time point when the subject experienced severe symptoms and TP3 is the time point when the subject was fully recovered.

**Table 2.** CDR3 Read counts from patient and healthy subject. Details regarding the ten highest CDR3 read counts recorded in each repertoire. For example, the most frequently occurring CDR3 sequence in Subject-A_TP2’s repertoire occurred 2,631 times in total. Subject-A_TP2 represents a patient repertoire while Subject-A_TP3 and 700010756 represent healthy repertoires. The read counts are sorted in descending order, and the table displays the first ten of them.

| Index | Read counts |  |  |
| --- | --- | --- | --- |
|  | Subject-A_TP2<br>(Patient) | Subject-A_TP3<br>(Healthy) | 700010756<br>(Healthy) |
| 0 | 2,631 | 979 | 9,901 |
| 1 | 1,777 | 375 | 215 |
| 2 | 1,459 | 700 | 153 |
| 3 | 1,236 | 475 | 102 |
| 4 | 1,177 | 457 | 97 |
| 5 | 909 | 416 | 80 |
| 6 | 908 | 386 | 70 |
| 7 | 899 | 377 | 64 |
| 8 | 811 | 340 | 63 |
| 9 | 716 | 319 | 62 |

Fig. 3b compares two repertoires from a single COVID-19 patient, Subject-A. Subject-A’s TP2 (severe symptoms) repertoire represents the patient repertoire, and TP3 (fully recovered) repertoire represents the healthy repertoire. Once again, we observed that the TP2 repertoire had more CDR3 sequences with significantly higher read counts than the healthy repertoire. The highest read count of the TP2 repertoire was 2,631, whereas that of the TP3 repertoire was 979 (Table 2). Both Fig. 3a and 3b highlight that the patient repertoires contain more CDR3 sequences with higher read counts.

### Visualization of individual sequences using t-SNE

Using the selected repertoires from OAS, we visualized each individual CDR3 sequence on a t-SNE chart [15]. We first extracted and embedded the ten most frequently occurring sequences in each repertoire (106 patient repertoires and 106 healthy repertoires), and simply projected each vectorized sequence in to a 2-dimensional vector space using t-SNE (Fig. 4). As shown in Fig. 4a, the sequences did not seem to have any clear patterns or form distinct clusters within the vector space.

**Fig. 4.**
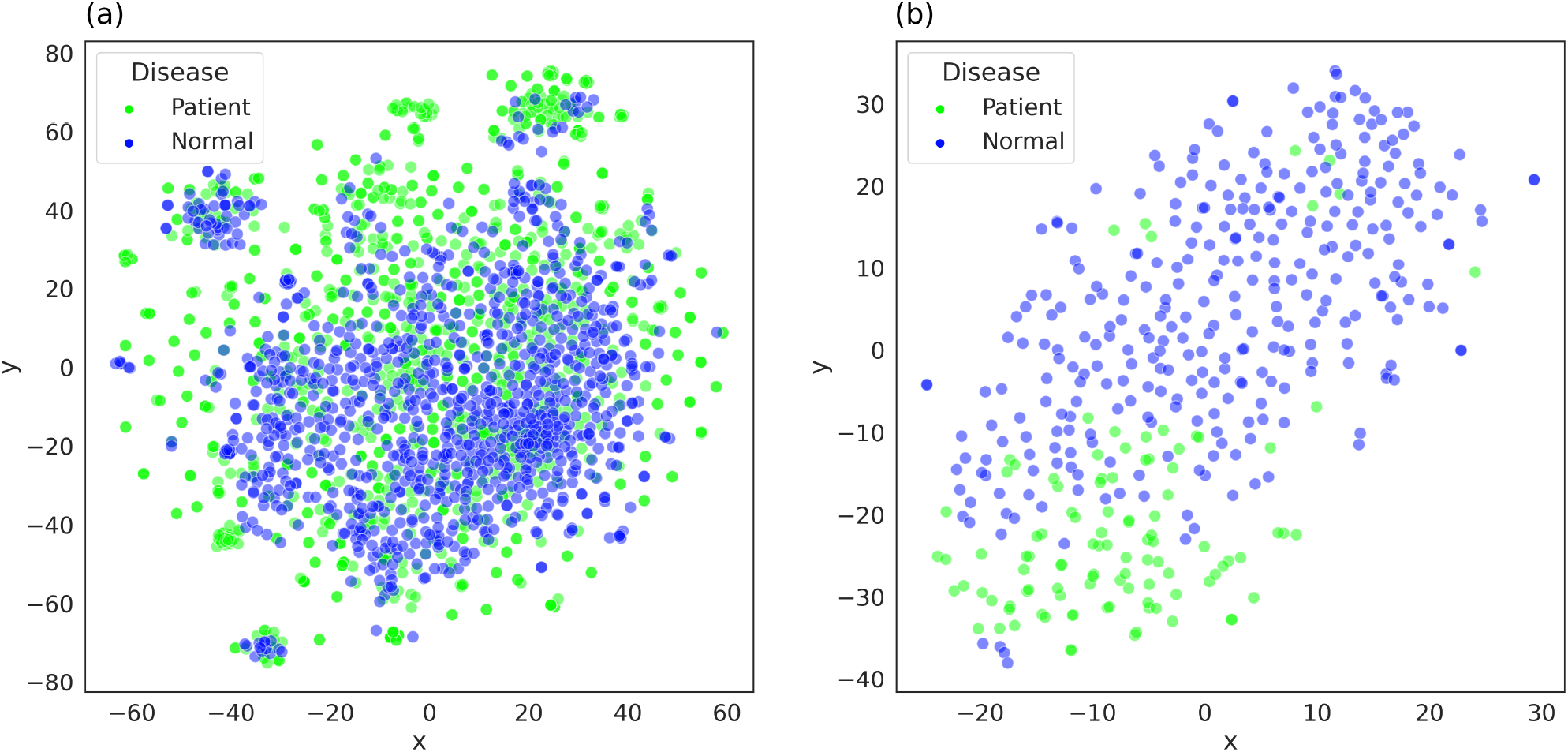
Visualization of embedded CDR3 sequences and BCR repertoires. **(a).** Result of embedding and visualizing individual CDR3 sequences from COVID-19 patients and healthy subjects using t-SNE. A total of 2,120 sequences (top 10 most frequent CDR3 sequences from 106 patients and 106 healthy subjects) were used. Every data point in the figure represents an individual CDR3 sequence—blue and green represent healthy subjects and COVID-19 patients, respectively. **(b).** Result of embedding and visualization of individual BCR repertoires of COVID-19 patients and healthy subjects using t-SNE. In both figures, a total of 428 repertoires (106 from COVID-19 patients and 322 from healthy subjects) were plotted. Details regarding specific datasets and studies used are summarized in Table S1.

### Visualization of individual repertoires using t-SNE

Rather than plotting individual sequences (Fig. 4), we decided to plot sets of sequences together, which, as have previously noted, represent BCR repertoires. We assumed that CDR3 sequences that occurred more frequently within a repertoire were more significant in the context of COVID-19 viral infection [16]. We thus sliced the 100 most frequently occurring CDR3 amino acid sequences from each repertoire and computed the sum of the vector representations of these sequences. This method allowed us to represent an entire repertoire as a single 100-dimensional vector. When we visualized the newly attained vector representations using t-SNE, we were able to successfully distinguish between 106 patient repertoires and 322 healthy repertoires (Fig. 4b).

As shown in Fig. 4b, we observed that the repertoires of COVID-19 patients mostly clustered in the bottom left corner, while those of healthy subjects clustered near the upper right region of the figure. There were, however, a few patient repertoires among the healthy repertoires. The positions of these repertoires and possible explanations will be discussed in a later section. Additionally, using the embedded representations of repertoires, we trained a binary classification model with XGBoost. The model achieved 87% mean accuracy in the classification of COVID-19 patients and healthy subjects given a 100-dimensional vector representation of a repertoire.

### Tracking the spatial and temporal movements of COVID-19 patient repertoires

Using the same public BCR dataset from OAS, we again sliced the 100 most frequently occurring CDR3 amino acid sequences from each repertoire to represent repertoires as 100-dimensional vectors. The t-SNE visualizations of these repertoire representations are shown in Fig. 5. We focused on tracking the spatial and temporal movements of specific repertoires from Kim’s study [7].

**Fig. 5.**
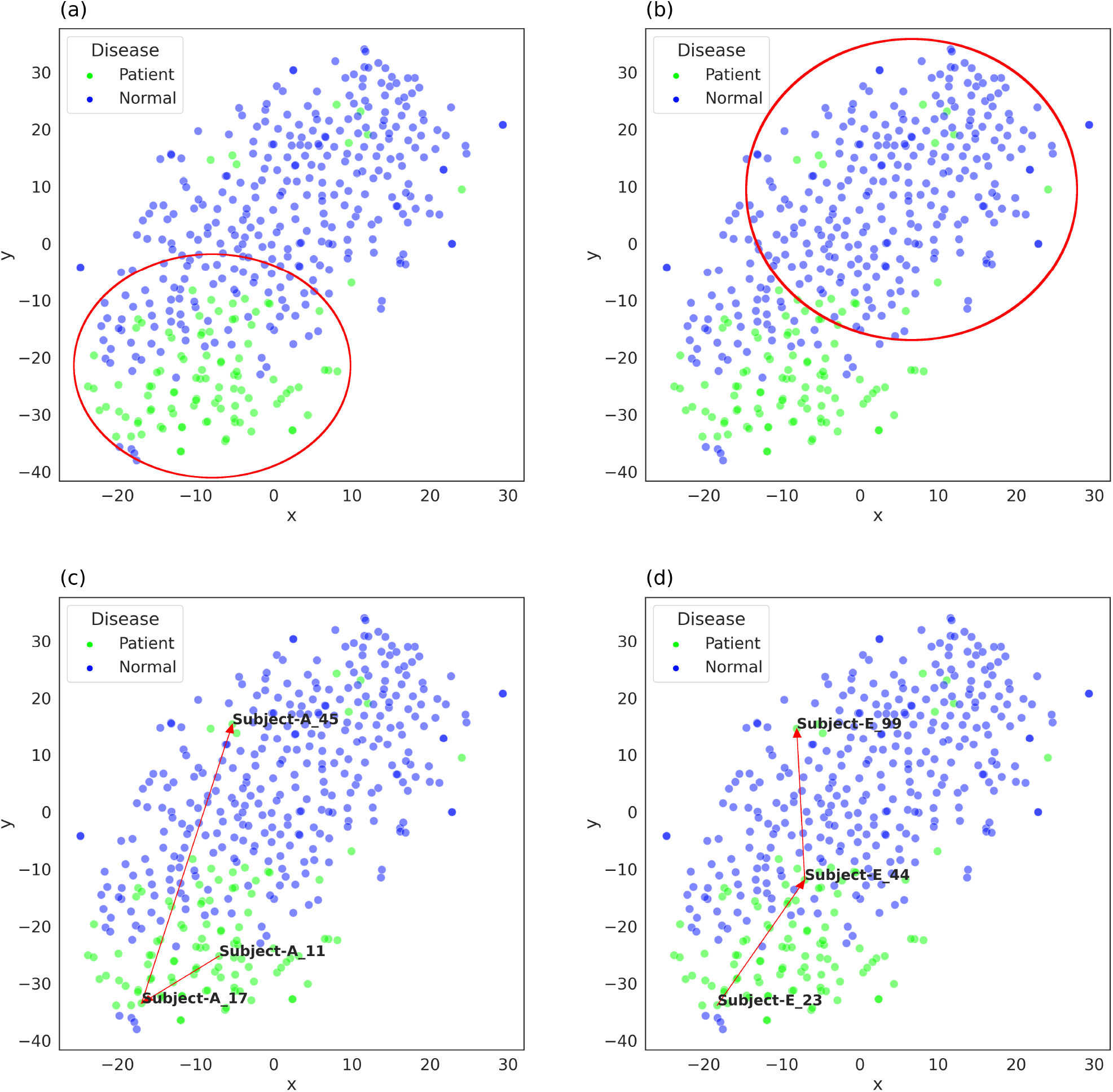
Visualization of embedded repertoires. Visualization of embedded repertoire representations of COVID-19 patients and healthy subjects from a study by Kim et al. **(a).** Repertoires of COVID-19 patients at 9 to 15 days after symptom onset formed their own cluster in the upper-left corner of the chart. The red circle indicates the disease cluster. **(b).** Repertoires of COVID-19 patients at 0 to 9 days and/or 21 to 45 days after symptom onset clustered in the bottom-right corner. The red circle indicates the healthy cluster. **(c).** Movement of Subject-A’s repertoires within the vector space over three time points (days 11, 17, and 45 after symptom onset). **(d).** Movement of Subject-E’s repertoires within the vector space over three time points (days 23, 44, and 99 after symptom onset).

We discovered that the repertoires could be largely categorized into two main groups—disease and healthy clusters. The disease cluster, as shown in Fig. 5a, is primarily composed of repertoires of COVID-19 patients at 9 to 15 days after symptom onset. The healthy cluster, as shown in Fig. 5b, mainly consists of repertoires of healthy subjects as well as patients at 0 to 9 days and/or 21 to 45 days after symptom onset.

For Subject-A and Subject-E, both of whom were COVID-19 patients, , we observed that their repertoires at three time points had remarkably interesting movements based on time, suggesting a potential association between our t-SNE charts and Kim’s chronological analysis of IGH repertoires of the two patients [7]. Kim highlighted that antibody clonotypes that were reactive against SARS-CoV-2 receptor-binding domains (RBDs) underwent swift class switching with minimal somatic mutations and divergence from the germ line. Isotypes and subtypes with low divergence thus suggest that the antibodies are actively reacting to viral invasion.

From Kim’s study, Subject-A’s IgG1 and IgG3 both had significantly low divergence on day 11 and even lower values on day 17 but ultimately had a large increase by day 45. These observations indicate that during a certain period of time, most likely around days 11 and 17, subject’s IgG1/3 antibodies have potentially bound to the RBDs of SARS-CoV-2 and neutralized the virus. An increased divergence on day 45 further suggests that the antibodies were no longer reacting as strongly as they did before. We observed a surprisingly similar pattern in our t-SNE chart (Fig. 5c). The repertoire of Subject-A was initially located near the center of the disease cluster on day 11 (A_TP1). By day 17 (A_TP2), the repertoire had moved closer to the disease cluster and away from the healthy cluster. However, by day 45 (A_TP3), the repertoire had moved to the other side, near the center of the healthy cluster.

Similarly, Kim’s study shows a gradual increase in divergence for Subject-E’s IgG1 and IgG3 across the three time points (day 23, 44, and 99). Over this time period, the reactivity of the subject’s IgG1/3 antibodies had potentially subsided after initially reacting strongly to the SARS-CoV-2 virus. Moreover, compared to Subject-A, Subject-E experienced a steadier and more gradual increase in divergence. These observations align well with the t-SNE visualization (Fig. 5d). The repertoire of Subject-E was inside the disease cluster on day 23 (E_TP1), but as time passed, the repertoire moved closer toward the healthy cluster (day 44; E_TP2) and eventually completely within the healthy cluster (day 99; E_TP3).

## Discussion

In this study, we introduced a novel computational approach of utilizing ProtVec, a deep-learning-based embedding technique for protein sequences, to represent a BCR repertoire as a 100-dimensional vector. Specifically, we discovered that by slicing the most frequently occurring CDR3 sequences and using the summation of vector representations of these sequences, we were able to not only effectively represent a subject’s entire BCR repertoire as a single data point within a vector space but also track the repertoire as the subject’s immunological status changed over time.

First, we assessed the efficacy of ProtVec to ensure that we could apply the embedding technique to represent BCR sequences in our study. As demonstrated in Fig. 2, the two t-SNE charts confirmed the effectiveness of ProtVec in representing different protein families. These selected protein families were grouped into respective clusters remarkably well, suggesting that ProtVec is more than capable of capturing unique features and characteristics of each protein family. It is especially important to note that ProtVec was also able to successfully distinguish protein families with similar mean amino acid sequence lengths (Fig. 2b). Moreover, we found that the embedded representations have other useful applications, such as supervised learning model training. The simple binary classification model we built using ProtVec representations was highly capable of accurately classifying protein families. Based on these results, we were confident that ProtVec would be an appropriate embedding method for our BCR repertoire analysis.

The distribution of frequencies of CDR3 amino acid sequences provided insight into the data we were analyzing (Fig. 3). A single BCR repertoire is typically composed of thousands, if not tens of thousands, of unique BCR amino acid sequences, each with its own frequency or read count [6]. The histogram of the CDR3 read counts revealed that while the vast majority of the sequences had low read counts, several sequences with abnormally high read counts were not uncommon. Fig. 3a compared repertoires from two different subjects, both in their 50s and with a similar number of unique BCR sequences. Fig. 3b, on the other hand, compared two repertoires from a single subject. Both figures ultimately revealed that patient repertoires, when compared to the healthy repertoires, had a much larger number of unique CDR3 sequences with significantly higher read counts.

We deduced that such differences can be associated with affinity maturation of B cells. Patients suffering from viral infection most often experience a rapid increase in the read counts of particular sequences due to the activation of B cells when BCRs bind to specific antigens [5]. Therefore, the CDR3 sequences on the right-hand sides of Fig. 3a and 3b were most likely the sequences that burgeoned due to of higher affinity to such antigens. We thus focused on small selections (the 100 most frequently occurring) of BCR sequences because we hypothesized that they were the sequences that had been rapidly increased by activation and hence possess much more significance in the comparison of COVID-19 patients and healthy subjects.

Our first attempt at using ProtVec to analyze BCR repertoires involved visualizing individual BCR amino acid sequences in a 2-dimensional vector space (Fig. 4). However, as Fig. 4 shows, the embedded sequences did not form any meaningful patterns or clusters that distinguished one class from another. There were small, indistinct clusters that included sequences from both patients and healthy subjects around the largest cluster; however, we deduced that their clustering was a result of other trivial external factors. We eventually arrived at the conclusion that we could possibly cluster or classify COVID-19 patients and healthy subjects when we analyzed only individual CDR3 amino acid sequences.

Subsequently, we represented entire repertoires by summing the vector representations of the 100 most frequently occurring CDR3 sequences (Fig. 5). The results highlight that this method can accurately distinguish between COVID-19 patients and healthy subjects. This discovery is especially meaningful because although we previously demonstrated that ProtVec could capture the unique characteristics of a BCR sequence (Fig. 2), there was no practical method for ProtVec to represent BCR repertoires themselves. Our new method of summing these vector representations, however, captures the unique features and characteristics of an entire repertoire, allowing us to visualize and analyze a repertoire in vector space. The ability to represent repertoires in low dimensions further enables us to track a subject’s repertoire based on time points.

As shown in Fig. 5a and 5b, plotting the repertoires of all the subjects at all the time points generates two main clusters (disease and healthy clusters). When we focused on two particular patients, Subject-A and Subject-E, we were able to track the movements of their repertoires over time (three time points). We discovered that the timeline and relative positions of subjects’ repertoires—whether they were located in the disease cluster or healthy cluster—aligned very well with the time points at which IGH clonotypes reactive against SARS-CoV-2 RBDs have been observed [7]. When the subject’s IgG1 and IgG3 had low divergence from the germ line, indicating that the antibodies were reacting to the virus, the repertoire was located within the disease cluster. On the other hand, when the subject’s IgG1 and IgG3 had relatively greater divergence, the repertoire was located within the healthy cluster. This result is immensely enlightening because it suggests that our repertoire representations do not just simply express a BCR repertoire in vector space; they can help in detecting protective immunity in a subject.

In conclusion, our novel approach enables researchers to computationally express and analyze a subject’s BCR repertoires within a vector space. Importantly, it allows us to pin-point one’s immunological status and track the repertoires spatial and temporal movement within a vector space as a patient undergoes treatment and recovery. We expect that our unconventional approach of utilizing protein embedding in repertoire analysis will allow the possibility for further BCR repertoire research in the near future.

## Author contributions

I.K. designed and proposed the research project. I.K., S.Y.B, S.K., and S.C. conducted data analysis. I.K., and S.Y.B. drafted the original manuscript. J.N., J.C., and B.G.K. discussed the research results together. All authors reviewed, edited, and approved the final draft.

## Acknowledgments

We thank Dongmin Kim for his excellent technical assistance. This research was supported by the K-BIO KIURI Program of the National Research Foundation funded by the Ministry of Science and ICT [NRF-2020M3H1A1073304].

## Data availability

The BCR repertoire representations can be accessed online via the following URL: https://data.mendeley.com/datasets/37tz3dkzkv/1

- 428 IGHG repertoire representations : 106 Covid-19 repertoire representations, 322 Healthy repertoire representations
- 311 IGHA repertoire representations : 23 Covid-19 repertoire representations, 288 Healthy repertoire representations

