## Supplemental Table 1 for "Computational analysis of B cell receptor repertoires in COVID-19 patients using deep embedded representations of protein sequences"

**Supplementary Information**

| Category | Author | Link |
|---|---|---|
| SARS-CoV-2 | Kim_2020 | https://stm.sciencemag.org/content/13/578/eabd6990 |
| SARS-CoV-2 | Montague_2020 | https://www.medrxiv.org/content/10.1101/2020.07.13.20153114v1.full.pdf |
| SARS-CoV-2 | Nielsen_2020 | https://www.researchsquare.com/article/rs-27220/v1%5D |
| Healthy | Bashford_2013 | https://genome.cshlp.org/content/23/11/1874 |
| Healthy | Bernat_2019 | https://www.frontiersin.org/articles/10.3389/fimmu.2019.00660/full |
| Healthy | Bonsignori_2016 | https://linkinghub.elsevier.com/retrieve/pii/S0092867416301246 |
| Healthy | Corcoran_2016 | https://www.nature.com/articles/ncomms13642 |
| Healthy | Eliyahu_2018 | https://www.frontiersin.org/articles/10.3389/fimmu.2018.03004/full |
| Healthy | Ellebedy_2016 | https://www.nature.com/articles/ni.3533 |
| Healthy | Galson_2015 | https://www.ncbi.nlm.nih.gov/pmc/articles/PMC4551417/ |
| Healthy | Galson_2016 | https://genomemedicine.biomedcentral.com/articles/10.1186/s13073-016-0322-z |
| Healthy | Ghraichy_2019 | https://www.biorxiv.org/content/10.1101/609651v1.full |
| Healthy | Montague_2020 | https://www.medrxiv.org/content/10.1101/2020.07.13.20153114v1.full.pdf |
| Healthy | Mroczek_2014 | https://www.frontiersin.org/articles/10.3389/fimmu.2014.00096/full |
| Healthy | Ohm-Laursen_2018 | https://www.frontiersin.org/articles/10.3389/fimmu.2018.01976/full |
| Healthy | Parameswaran_2013 | https://www.ncbi.nlm.nih.gov/pmc/articles/PMC4136508/ |
| Healthy | Roskin_2020 | https://www.nature.com/articles/s41590-019-0581-0 |
| Healthy | Rubelt_2016 | https://www.nature.com/articles/ncomms11112 |
| Healthy | Schanz_2014 | https://journals.plos.org/plosone/article/comments?id=10.1371/journal.pone.0111726 |
| Healthy | Schultheiss_2020 | https://www.sciencedirect.com/science/article/pii/S107476132030279X |
| Healthy | Sheng_2017 | https://www.frontiersin.org/articles/10.3389/fimmu.2017.00537/ |
| Healthy | Simonich_2019 | https://www.nature.com/articles/s41467-019-09481-7 |
| Healthy | Turchaninova_2016 | https://www.nature.com/articles/nprot.2016.093 |
| Healthy | Vander_Heiden_2017 | https://pubmed.ncbi.nlm.nih.gov/28087666/ |
| Healthy | Vergani_2017 | https://www.frontiersin.org/articles/10.3389/fimmu.2017.01157/full |
| Healthy | Waltari_2018 | https://www.frontiersin.org/articles/10.3389/fimmu.2018.00628/full |

**Table S1. Authors and links to their studies.** These data, which are available at OAS, were used as datasets for our study. The table includes details regarding disease category (SARS-CoV-2 and healthy), author names, and links to their studies. For the attributes used to search the OAS sequences, we set ‘Chain’ to ‘Heavy’, ‘Isotype’ to ‘IGHG’, ‘Disease’ to ‘SARS-COV-2’ or ‘None’, ‘BSource’ to ‘PBMC’, ‘Vaccine’ to ‘None’, and ‘Species’ to ‘human’.
